# ASO-mediated mRNA silencing enables functional analysis and selective depletion of the human microbiota *Prevotellaceae*

**DOI:** 10.64898/2026.04.17.719208

**Authors:** Valentina Cosi, Vincent Lau, Petia Kovatcheva-Datchary, Youssef El Mouali, Toby Wilkinson, Victoria Gebler, Linda Popella, Franziska Faber, Till Strowig, Jörg Vogel

## Abstract

The *Prevotellaceae* family comprises abundant, taxonomically diverse bacteria of the human microbiota that exhibit remarkable intraspecies variability and distinct phenotypes during host-microbe interactions. Yet functional investigations are hampered by the limited genetic tractability of this medically important bacterial family, rendering it understudied. Here, we apply antisense oligomer (ASO) technology to selectively inhibit translation of single or multiple mRNAs across nine *Prevotellaceae* species. Using *ftsZ-* and *mreB-*related phenotypes as morphogenic readouts, we demonstrate that ASOs can function as a tunable system to study essential gene function. Further, we show that ASOs can selectively deplete target species from a synthetic Bacteroidales community. These results establish ASOs as practical tools for functional genomics and community modulation in otherwise genetically intractable anaerobic microbiota members.

## INTRODUCTION

Microbiome research is a rapidly advancing field that drives the development of alternative strategies for health promotion and disease treatment (Hou et al. 2022). Major objectives are the elimination of pathogens or pathobionts while preserving overall microbiome structure, and the precise functional modulation of individual members within complex microbial communities. *Prevotellaceae* are a prominent example of a keystone microbiota family, known for their high prevalence in livestock and their beneficial roles in ruminants (Kou et al. 2024). Distinct *Prevotellaceae* species also colonize multiple sites of the human body, such as the oral cavity, skin, genital tract, and gastrointestinal tract (Tett et al. 2021).

The *Prevotellaceae* family includes *Segatella copri*, formerly known as *Prevotella copri*, which is a highly abundant human gut commensal and the most abundant species in 34% of individuals (Pasolli et al. 2017; Tett et al. 2019). Abundance of *Segatella* has been linked with improved metabolic health (Kovatcheva-Datchary et al. 2015; Haro et al. 2017; Marungruang et al. 2018; Roager et al. 2019), but also with specific diseases such as hypertension, insulin resistance, and inflammatory conditions (Scher et al. 2013; Pedersen et al. 2016; Dillon et al. 2016; Li et al. 2017). Importantly, investigations into a suspected role of *S. copri* in rheumatic disorders revealed striking differences in the immunostimulatory and arthritis-induction potential of different *Segatella* strains (Amend et al. 2023; Nii et al. 2023; Maeda et al. 2026). Further metagenomic investigations of newly isolated strains revealed that *S. copri* constitutes a broader taxonomic complex of 13 distinct species that are hard to distinguish at the 16S rRNA sequence level (Tett et al. 2019; Blanco-Míguez et al. 2023).

The fact that different members of the *Segatella copri* complex can coexist in individual human microbiomes has created a need for general functional genomic tools to understand the observed diversification across phenotypes, metabolic preferences, antibiotic susceptibility, and host cell- and microbiome-interactions (Blanco-Míguez et al. 2023; Xiao et al. 2024). However, robust plasmid transformation and chromosomal gene disruption only work in select strains of *Segatella* spp. (Li et al. 2021). To overcome these limitations and accelerate gene-function analysis within the *Prevotellaceae* family, we adopt an alternative approach to post-transcriptional control that employs cellular delivery of short antisense oligomers (ASOs) for programmable silencing of target mRNAs of interest.

Antimicrobial ASOs have proven a viable non-genetic tool for selective gene repression, without the need for auxotrophies or antibiotic selection markers (Vogel et al. 2025; Moustafa et al. 2025). Typically designed to sequester the ribosome-binding site (RBS) of the target mRNA of interest and delivered via a cell-penetrating peptide (Good and Nielsen 1998), these agents have primarily been used to kill pathogens by blocking the synthesis of essential proteins (El-Fateh et al. 2024; Moustafa et al. 2025). Going beyond this, recent studies have leveraged ASOs to address enzyme functions in a difficult-to-transform environmental bacterium (Yokoi et al. 2022), to decrease species abundance in a small synthetic community composed of four physiologically unrelated bacteria (Hizume et al. 2024), or to perform functional genomics in diverse bacteriophages (Gerovac et al. 2025). Nonetheless, while ASOs have been used with success in taxonomically diverse aerobic bacteria, attempts in strictly anaerobic bacteria have so far failed to achieve selective mRNA repression (Barbosa et al. 2015; Cosi et al. 2025). In this work, we utilize ASO technology to selectively inhibit mRNA translation in numerous strictly anaerobic members of the *S. copri* complex and other members of the *Prevotellaceae*. Using ASO multiplexing, we demonstrate simultaneous mRNA targeting of up to three different genes, breaking down the barrier for gene-function studies in genetically intractable *Prevotellaceae*. Furthermore, by demonstrating the selective depletion of an *S. copri* strain from a physiologically relevant gut microbiota community, we provide proof-of-concept for ASOs as a promising research tool for unraveling interspecies interactions among abundant members of the human microbiome.

## RESULTS

### ASOs targeting the essential gene acpP kill the Segatella copri type strain

Given the documented limited efficacy of ASOs in several anaerobic bacteria (El-Fateh et al. 2024; Cosi et al. 2025), it was unclear whether ASOs would work in *Prevotellaceae*, whose members generally require strict anaerobic conditions for handling and cultivation. To obtain a proof-of-concept for successful ASO delivery via CPPs and subsequent mRNA inhibition (Fig. 1a), we designed an 11-mer ASO to sequester by antisense pairing the RBS of *acpP* mRNA in the *S. copri* type strain DSM 18205⍰ (Fig. 1b). The *acpP* mRNA, which encodes the essential acyl carrier protein for fatty acid biosynthesis, has been the most popular target of antimicrobial ASOs (Moustafa et al. 2025; El-Fateh et al. 2024). As control, we used a scrambled sequence of the *acpP* ASO (termed scr ASO), with no predicted match in the RBS of any other mRNA based on the DSM 18205^T^ genome annotation (Fig. 1b). We chose peptide nucleic acid (PNA) as ASO chemistry, as this modality shows high efficacy in mRNA inhibition (Ghosh et al. 2024) and is resistant to nucleases (Moreira et al. 2024).

**Fig. 1:**
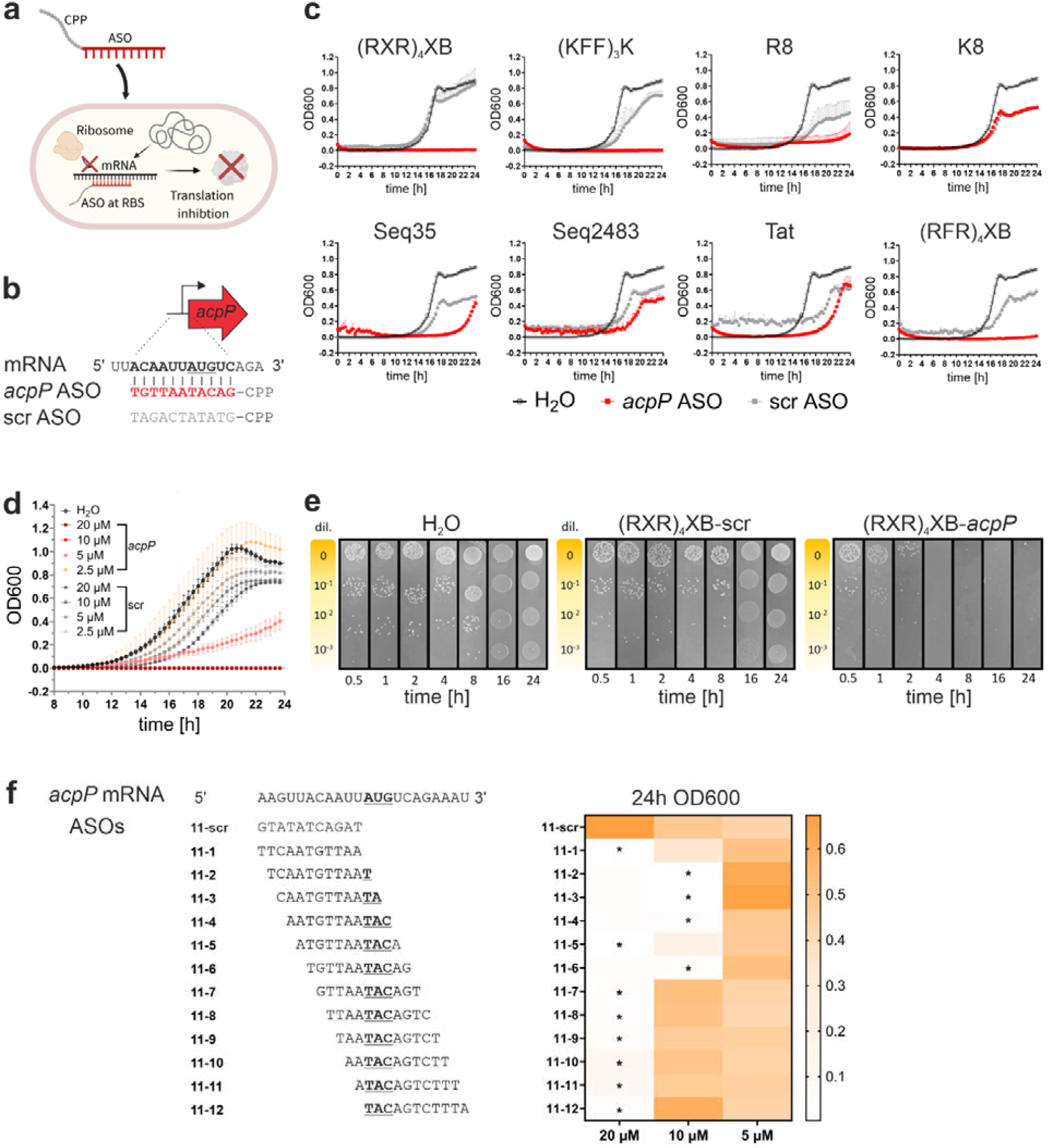
CPP-ASOs targeting the essential gene acpP demonstrate bactericidal effect in Segatella copri DSM 18205^T^. **a**, ASO mode of action: Following uptake into bacterial cells via CPP-mediated delivery, ASOs accumulate in the cytoplasm and inhibit the translation of target mRNAs via binding to the RBS. Illustration created with BioRender. **b**, CPP-ASO design for *S. copri* DSM 18205^T^ *acpP*. A scrambled sequence control with no complementarity to any translation start codon within the *S. copri* genome is shown below. **c**, Growth kinetics of *S. copri* treated with different 20 µM CPPs conjugated to the *acpP*-ASO. Growth curves are depicted as the mean OD600 of three independent starting cultures, with standard deviations shown. **d**, Minimum inhibitory concentration (MIC, OD600 <0.1) of (RXR)_4_XB-*acpP* in *S. copri*. Data are shown as the mean of two independent experiments; error bars indicate standard deviation. **e**, Spotting assay of SC treated with H_2_O, (RXR)_4_XB-scr or (RXR)_4_XB-*acpP*. A representative image of three independent starting cultures is shown. **f**, (Top) ASO sequences tiling the *S. copri acpP* mRNA translation initiation region. The scrambled sequence of ASO 11-6 served as control. (Bottom) OD600 value is shown in a white to orange color gradient; MIC value is indicated with an asterisk. Data are based on two independent experiments; median values are shown.

Screening eight different CPPs for their ability to deliver the *acpP*-ASO into DSM 18205⍰ (Table 1), we concluded that (RXR)_4_XB and (KFF)_3_K were the most effective delivery vehicles, achieving complete growth inhibition at 20 µM. CPPs R8, Seq335, and (RFR)_4_XB, when coupled to the *acpP*-ASO, also showed initial growth inhibition, yet bacterial outgrowth occurred within 24 hours. CPPs K8, Seq2483, and Tat caused unspecific growth inhibition, as inferred from the scrambled control (Fig. 1c). When assessing potential unspecific cytotoxic effects of the two best CPPs in their unconjugated form, (RXR)_4_XB showed no antimicrobial activity up to 40 µM, whereas unconjugated (KFF)_3_K impaired growth at 20 µM (Extended Data Fig. 1a). Consequently, we chose (RXR)_4_XB for ASO delivery in all subsequent experiments.

**Table 1:**
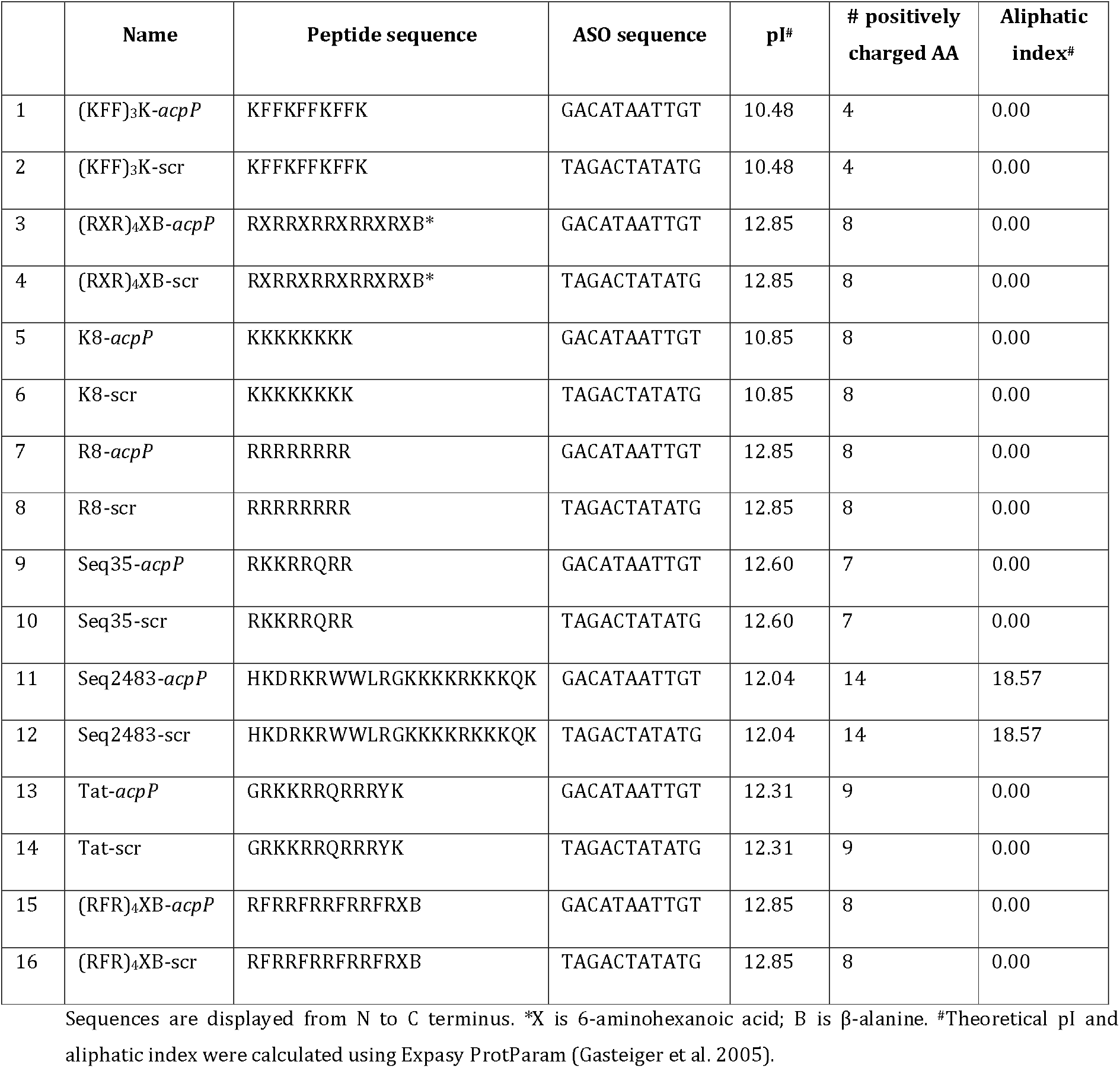
Cell-penetrating peptide conjugated ASOs for acpP screen in S. copri DSM 18205^T^.

The minimum inhibitory concentration (MIC) of the (RXR)_4_XB-*acpP* ASO, measured in a broth dilution assay with the strain DSM 18205⍰, was determined as 10 µM (Fig. 1d). To distinguish between a bacteriostatic or a bactericidal effect, we performed quantification of agar-based colony-forming units (CFUs) upon ASO treatment. Since the minimal bactericidal concentration is usually higher than the MIC (Bury-Moné 2014), we used 2x MIC to determine bactericidal effects. Spotting assays demonstrated bactericidal activity at 20 µM, with no recovery of CFUs after 4 h (Fig. 1e). These results show that ASOs delivered by the CPP (RXR)_4_XB targeting the essential gene *acpP* are potent bactericidal agents against *S. copri*.

ASO potency depends on multiple parameters, including uptake efficacy, target region, hybridization strength with the cognate mRNA target, number of off-targets, nucleobase composition, and secondary structure formation (Jung et al. 2023). To probe the optimal ASO-targeting site, we tiled the *acpP* mRNA in the -11 to +11 nucleotide window relative to the AUG start codon with twelve different 11-mer ASOs. We observed effective inhibition at 10 µM in the region from -10 to +5, except for ASO 11_5 (-7 to +4). Outside this region, the MIC increased by two-fold (Fig. 1f).

Using the data from our tiling assay, we investigated whether any ASO parameters, such as melting temperature, self-complementarity, GC content, or off-target number, predict ASO potency. The top-performing ASOs (MIC 10 µM) have predicted melting temperatures between 33.8 and 43.2°C according to MASON (Extended Data Fig. 1b), well within the range reported for PNA:mRNA melting temperatures of successful antimicrobial ASOs (Goltermann et al. 2019). However, we found that melting temperature alone is an insufficient predictor for ASO potency. For example, ASO 11_9 has the highest predicted melting temperature (47.1°C) of all the tiling ASOs, but not the lowest MIC. Hence, we also examined the number of self-complementary bases and purine content, both of which are important for the formation of secondary structures and solubility of ASOs, which can influence the efficacy of translation inhibition (Jung et al. 2023). Our best ASOs neither have the lowest number of self-complementary bases nor the highest purine content, suggesting that no clear association exists between these factors and the observed MIC (Extended Data Fig. 1b).

Since a high off-target number might decrease the amount of available ASO for the intended target, similar to the “sponging” effect described for sRNAs (Papenfort and Melamed 2023), we also predicted the number of potential ASO off-target sites in all *S. copri* DSM 18205^T^ genes. While the predicted off-target numbers ranged from 122 for ASO 11_9 and ASO 11_10 to 259 for ASO 11_12, we found no clear correlation with the respective MIC (Extended Data Fig. 1b). Since multiple factors seem to determine ASO efficacy, it is advisable to design and test several different ASOs targeting the mRNA region from -10 to +5 relative to the AUG start codon to identify the most potent candidate.

### ASOs have broad applicability within the Prevotellaceae family

To assess ASO susceptibility across the *S. copri* complex, we first tested *acpP* ASOs in ten additional strains from seven different *Segatella* species. The *acpP* site was conserved in eight of these strains, but not in *Segatella brunsvicensis* NI025 and *Segatella hominis* HDD12 (Fig. 2a). Notably, within the *S. copri* complex, these latter two species also share the lowest average nucleotide identity with *S. copri* DSM 18205^T^ (Blanco-Míguez et al. 2023). Eight out of the tested ten strains were found to be susceptible to (RXR)_4_XB-*acpP* within the concentration range used above for the type strain (20 to 2.5 µM), with the best MIC value as low as 5 µM (*S. copri* RPC01; Fig. 2a). We observed no growth inhibition for strains *S. copri* HDC01 and *S. hominis* HDD12. Since the *acpP*-ASO led to a moderate growth delay in the strain HDC01 at 20 µM, we increased the ASO concentration to 40 µM. At this concentration, HDC01 growth was fully inhibited, but the scrambled control also impaired growth (Extended Data Fig. 1c). As the unconjugated CPP (RXR)_4_XB also started to show growth-inhibiting effects at 40 µM in *S. copri* DSM 18205^T^ (Extended Data Fig. 1a), the CPP-ASO conjugates likely exhibit intrinsic cytotoxicity at this concentration. Based on these results, we conclude that four additional human isolate *S. copri* strains and six additional *Segatella* species, except *S. hominis* HDD12, demonstrate growth inhibition ≤40 µM upon treatment with an (RXR)_4_XB-ASO targeting *acpP*.

**Fig. 2:**
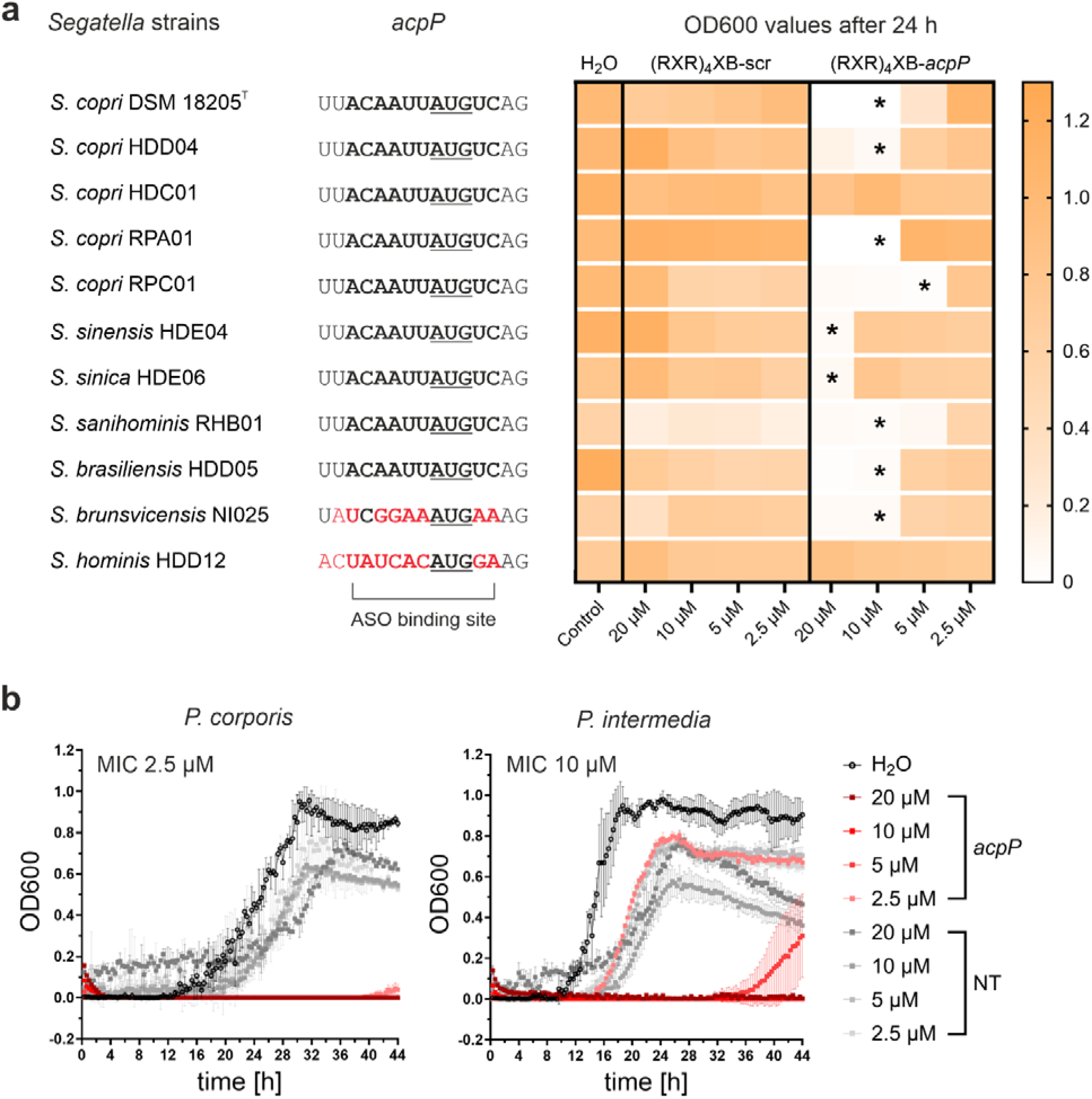
CPP-ASOs allow for gene-specific modulation in 13 strains of the Prevotellaceae family. **a**, MIC (OD600 <0.1) determination of twelve strains of the *S. copri* complex. *acpP* mRNA sequences of *Segatella copri* complex strains (ScC). ASO binding site shown in bold, base differences with respect to SC shown in red. OD600 values after 24 h depicted in a white to orange color gradient shown for individual strains treated with H_2_O, titration series of the respective (RXR)_4_XB-*acpP* or (RXR)_4_XB-scr; MIC value is indicated with an asterisk. Data are based on two independent experiments; median values are shown. **b**, MIC (OD600 <0.1) determination of (RXR)_4_XB-*acpP* for *P. corporis* DSM 18810^T^ and *P. intermedia* DSM 20706^T^ together with H_2_O and (RXR)_4_XB-non-targeting controls. Data are shown as the mean of three independent experiments; error bars indicate standard deviation.

To advance ASOs as a tool for the whole *Prevotellaceae* family, we also targeted the *acpP* mRNAs in the commensal cervical strain *Prevotella corporis* DSM 18810^T^ and the oral pathogenic strain *Prevotella intermedia* DSM 20706^T^. Of the two, *P. corporis* appeared more sensitive to ASOs with an MIC of 2.5 µM, while *P. intermedia* showed an MIC of 10 µM (Fig. 2b). However, we did observe slight unspecific growth defects upon treatment with the scrambled ASOs, suggesting that these *Prevotella* strains are generally more sensitive to (RXR)_4_XB-conjugated ASOs than *S. copri*. Overall, our results demonstrate that ASOs can serve as a programmable tool for translation inhibition across multiple genera within the *Prevotellaceae* family.

### ASOs targeting ftsZ and mreB mRNAs induce expected morphological phenotypes

Many *S. copri* complex strains are difficult, if not impossible, to modify by plasmid-based genetic tools (Li et al. 2021). Thus, to demonstrate the applicability of ASOs for studying gene functions in these species, we targeted mRNAs with predictable phenotypes as direct readouts. We designed ASOs to inhibit the mRNAs of the cell division Z-ring protein FtsZ or the actin-like murein formation protein MreB (Fig. 3a). Both genes are essential in many bacterial species, and studies using small molecule inhibitors or inducible gene depletion showed altered cell morphology prior to complete growth arrest (Soufo and Graumann 2003; Gitai et al. 2004; Kruse et al. 2005; Paradis-Bleau et al. 2007). We selected different *Segatella* species: *S. copri* HDC01, *S. copri* RPC01, *Segatella sinica* HDE06, and *Segatella brasiliensis* HDD05, because of the lack of robust genetic tools for these species and the observed low to undetectable DNA uptake via conjugation (Li et al. 2021).

**Fig. 3:**
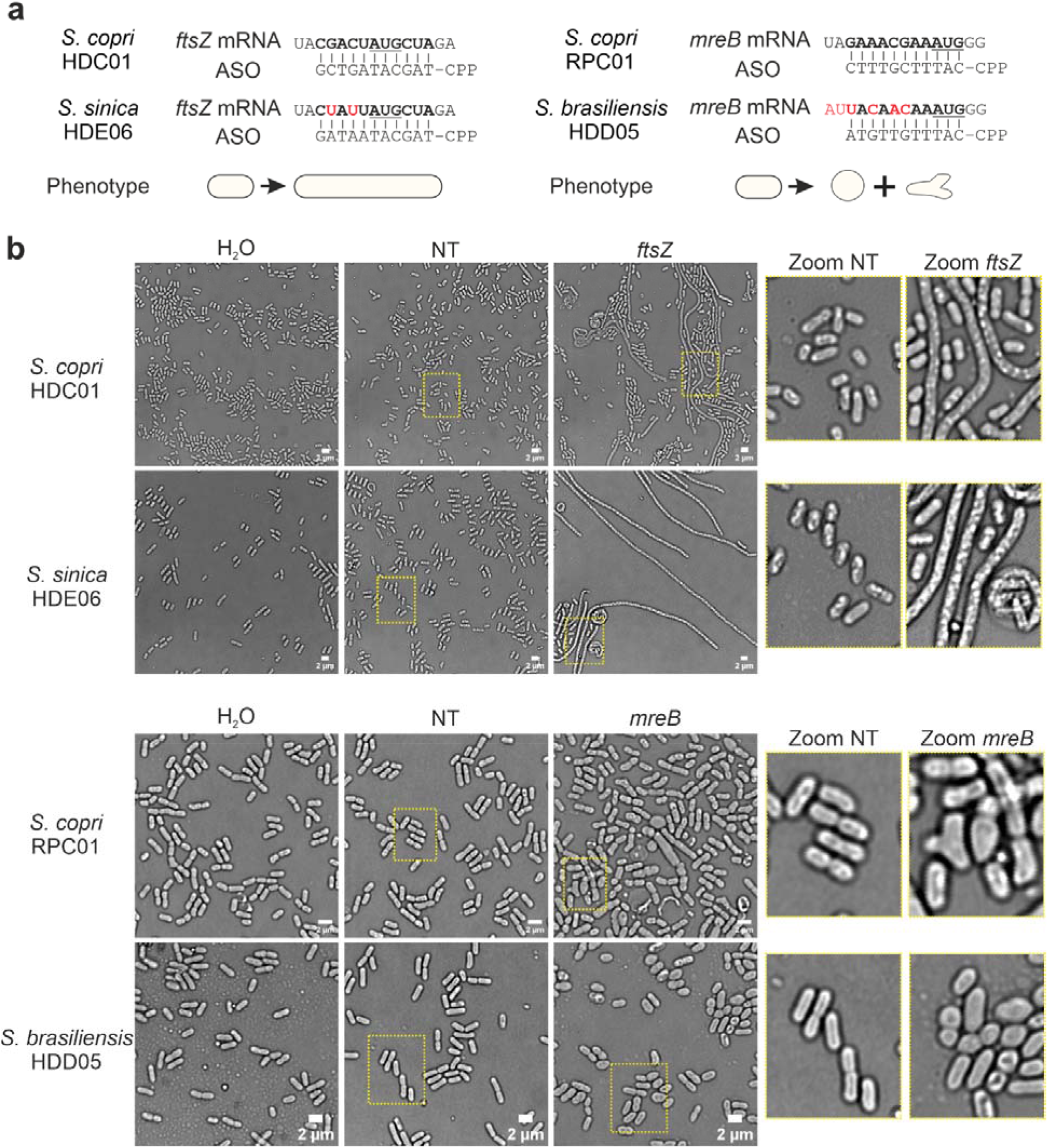
Phenotype-based readout allows to investigate ASO efficiency in genetically intractable Segatella strains. **a**, mRNA sequences of *ftsZ* from *S. copri* HDC01 and *S. sinica* HDE06 (left), as well as *mreB* from *S. copri* RPC01 and *S. brasiliensis* HDD05 (right). ASO binding site shown in bold, base differences with respect to SC highlighted in red. Cognate ASO sequences are shown below their respective mRNA targets, together with the expected phenotypes depicted below. Illustration created with BioRender. **b**, Representative bright field images from two independent experiments of *S. copri* HDC01 and *S. sinica* HDE06 treated with (RXR)_4_XB-ASOs targeting the *ftsZ* mRNA (top) or *S. copri* RPC01 and *S. brasiliensis* HDD05 treated with (RXR)_4_XB-ASOs targeting the *mreB* mRNA (bottom), as well as H_2_O and ASO non-targeting control (NT) for each strain. Scale bar, 2 µm.

Targeting the *ftsZ* mRNAs in *Segatella* species, we readily observed the expected elongated cell phenotype and filamentation, with cell lengths of up to 40 µm, consistent with the current model that FtsZ depletion impairs Z-ring formation (Sundararajan et al. 2015). Similarly, targeting the *mreB* mRNA led to rounded and dysmorphic cell shapes with sporadic formation of an additional cell pole (Fig. 3b), which is consistent with the uneven, rounded, and often biforked cell morphology observed upon MreB depletion in other species (Strahl et al. 2014; Kawazura et al. 2017). Importantly, these ASO-induced phenotypes were dose-dependent, enabling investigation of essential gene knockdown-induced morphologies below MIC (Extended Data Fig. 2a, 2b, and 2c). Collectively, these data demonstrate that ASOs enable phenotype-based functional genomic studies in genetically intractable strains of *Segatella*.

### Simultaneous ASO treatment for multiplex mRNA inhibition and gene function studies

As shown above, the *acpP* mRNA proved to be a robust target across all tested *Segatella* strains, with the striking exception of *S. hominis* HDD12, which showed no detectable growth inhibition. Targeted resequencing of the *acpP* site in HDD12 ruled out a spontaneous mutation in the genome that might compromise ASO binding. A search for alternative explanations revealed that the HDD12 genome carried two additional *acpP* gene candidates, both with different RBS-sequences from *acpP_1* (first gene to be annotated as *acpP* in the NCBI genome file CP137559.1), which we initially targeted above for MIC determination (Fig. 4a). All three predicted AcpP proteins showed high similarity to one another, including helical secondary structures, and a conserved serine residue motif important for 4’-phosphopantetheine transfer, all of which are necessary for the biological activity of AcpP in *E. coli* (Zhu and Cronan 2015) (Fig. 4b). Consequently, we sought to determine whether ASO-mediated killing of *S. hominis* HDD12 required a simultaneous inhibition of all three *acpP* mRNAs proteins. Using cognate ASOs for each of the three mRNAs (Fig. 4c), we observed no growth inhibition with single ASOs targeting either *acpP*_1 or *acpP*_2. Targeting *acpP*_3 alone resulted in a growth delay at 20 µM (Extended Data Fig. 3a). In contrast, simultaneous targeting of all three *acpP* mRNAs impaired growth at a combined ASO concentration of 2.5 µM and killed this strain at a total ASO concentration of 20 µM, similar to the *Segatella* strains with only one gene copy (Fig. 4d and Extended Data Fig. 3a). These results suggests that *S. hominis* HDD12 harbors three *acpP* gene copies that encode functional redundant AcpP proteins, as simultaneous inhibition of all three mRNAs was required for effective growth inhibition.

**Fig. 4:**
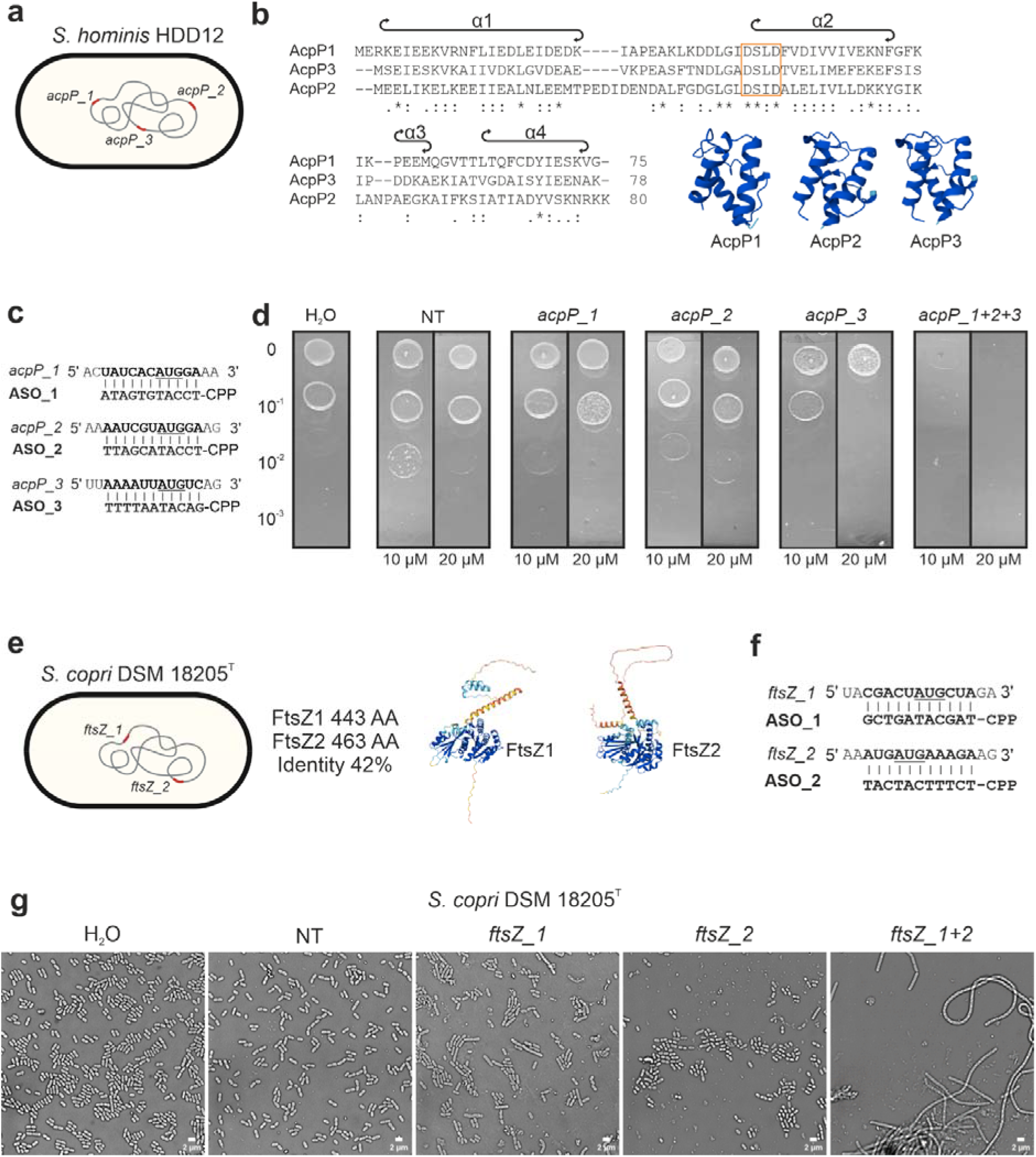
Multiplex application of CPP-ASOs enables investigation of gene duplication and functionality in members of the S. copri complex. **a**, The *S. hominis* HDD12 genome encodes three *acpP* copies. Illustration created with BioRender **b**, Sequence alignment of AcpP1, AcpP2, and AcpP3 created with Clustal Omega, together with modeled protein structures by AlphaFold 3. Alpha helix regions are indicated above the alignments, and the conserved serine motif is highlighted by an orange box. **c**, mRNA sequences of *acpP_1, acpP_2*, and *acpP_3* with respective ASO sequences. **d**, Spotting assay of *S. hominis* HDD12 upon treatment with the indicated ASOs. A representative image of three independent starting cultures is shown. **e**, *S. copri* DSM 18205^T^ genome with two *ftsZ* copies (left). Illustration created with BioRender. FtsZ protein identity was determined using the EMBOSS NEEDLE pairwise sequence alignment tool. The three-dimensional structure of FtsZ proteins was modeled by AlphaFold (right). **f**, mRNA sequences of *ftsZ_1* and *ftsZ_2* with respective ASO sequences. **g**, Representative bright field images of *S. copri* DSM 18205^T^ from two independent experiments, treated with H_2_O, 10 µM non-targeting control (NT), or *ftsZ* (RXR)_4_XB-ASOs. Scale bar, 2 µm.

Interestingly, when we tested *ftsZ* and *mreB* inhibition in *S. copri* DSM 18205^T^, we observed the expected change in morphology for *mreB*, but not for *ftsZ* (Extended Data Fig. 3b). While multiple *acpP* genes have been described in different bacterial species (Chen et al. 2018; Zhu et al. 2019; Yin et al. 2022), multiple *ftsZ* genes have only been reported in archaeal species (Liao et al. 2021; Santana-Molina et al. 2023). However, a closer inspection of the *S. copri* DSM 18205^T^ genome revealed two *ftsZ* genes (Fig. 4e). The encoded proteins both showed high amino acid sequence similarity to *E. coli* FtsZ and contained structural elements (a conserved globular core responsible for GTP binding and a C-terminal binding region for membrane anchoring proteins) that are necessary for FtsZ function (Extended Data Fig. 3c) (McQuillen and Xiao 2020). To test whether both proteins acted redundantly in Z-ring formation, we used ASOs to target their mRNAs alone or in combination (Fig. 4f). Only the simultaneous targeting of both *ftsZ* mRNAs induced cell elongation, while inhibition of one or the other mRNA alone had a minor effect (Fig. 4g). Taken together, our ASO approach suggests that *S. copri* DSM 18205^T^ encodes two functional redundant FtsZ proteins.

Next, we employed ASO multiplexing to investigate FtsZ duplication in other *Segatella* species. *Segatella sanihominis* RHB01 also contains two *ftsZ* copies, but while *ftsZ_1* is very similar to *ftsZ_1* of *S. copri* DSM 18205^T^ (92% protein identity), *ftsZ_2* is smaller and lacks parts of the GTP-binding globular domain as well as C-terminal structural elements that are critical for polymerization and interaction with other proteins involved in cell division in *E. coli* (Extended Data Fig. 4a) (Radler and Loose 2024). Microscopy images of *S. sanihominis* RHB01 treated with ASOs reveal that *ftsZ_1* inhibition alone leads to cell elongation similar to *S. copri* HDC01 and *S. sinica* HDE06 (Fig 3b and Extended Data Fig. 4b), but inhibition of *ftsZ_2* alters cell morphology without causing drastic elongation. Multiplex inhibition of *ftsZ1* and *ftsZ2* did not further increase cell elongation compared to single *ftsZ_1* inhibition (Extended Data Fig. 4b), suggesting that silencing of *ftsZ_1* alone is sufficient to inhibit Z-ring formation and subsequent cell division. FtsZ2 appears to play a role in cell morphology, but its exact function remains unknown. Taken together, these results show that the parallel application of ASOs reveals distinct functions of FtsZ1 and FtsZ2 in *Segatella* strains.

### Selective removal of Segatella copri from a Bacteroidales community

For ASOs to be suitable tools for microbiota modulation, they must demonstrate high species selectivity, ensuring precise targeting of the intended organism while leaving other commensal bacteria unaffected. Therefore, we investigated whether an ASO targeting the *S. copri* DSM 18205^T^ *acpP* affects other members of the order Bacteroidales, namely *Bacteroides ovatus* DSM 1896^T^ and *Bacteroides thetaiotaomicron* DSM 2079^T^. We chose these species because they are phylogenetically close to *Segatella* and occupy a similar gut niche (Adak and Khan 2018; Culp and Goodman 2023). The *acpP* RBS sequences differ among these three species, and the cognate *S. copri* ASO contains two mismatches within the corresponding *B. ovatus* and *B. thetaiotaomicron acpP* binding region. These mismatches should prevent ASO-mediated repression of the respective *Bacteroides* mRNAs, as in previous studies, where two mismatches within non-terminal ASO-binding positions abrogated translation inhibition (Jung et al. 2023) (Fig. 5a, top left). In addition, a transposon-insertion screen suggested that *acpP* is not an essential gene in *B. thetaiotaomicron* (Liu et al. 2021). Indeed, in preliminary monoculture testing, the *S. copri* (RXR)_4_XB-*acpP* ASO did not impair the growth of *B. ovatus* or *B. thetaiotaomicron* (Fig. 5a).

**Fig. 5:**
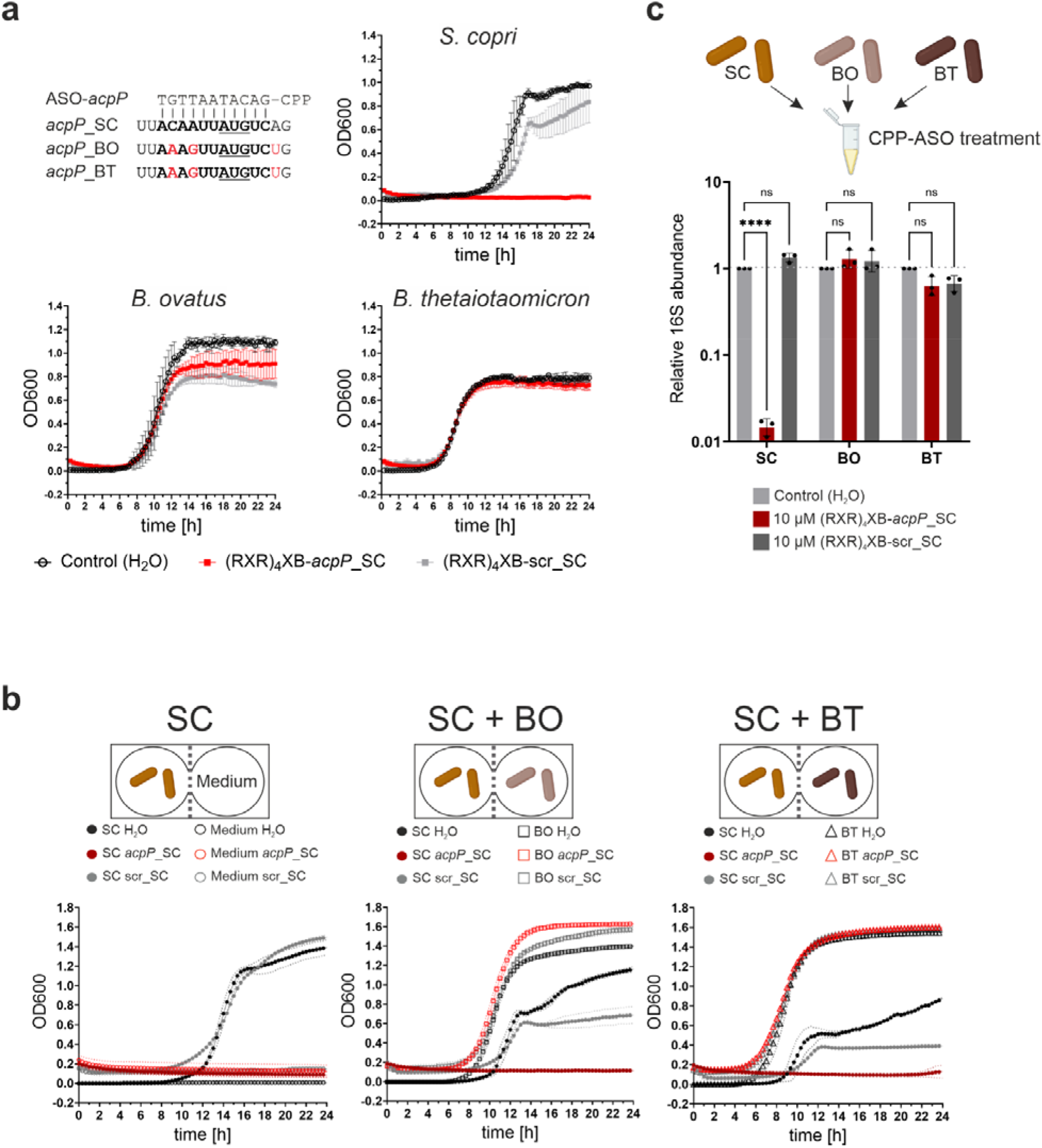
Selective modulation of a Bacteroidales community using ASOs. **a**, (Top left) Alignment of the *acpP* ASO and *acpP* mRNAs of *S. copri* DSM 18205^T^ (SC), *B. ovatus* DSM 1896^T^ (BO), and *B. thetaiotaomicron* DSM 2079^T^ (BT), the binding site is shown in bold, and base differences with respect to SC are shown in red. (Bottom left, right) Growth curves of SC, BO, and BT in monoculture upon treatment with H_2_O, 10µM (RXR)_4_XB-scr or SC (RXR)_4_XB-*acpP*. Data are shown as the mean of three independent starting cultures with standard deviation. **b**, Growth curves from co-culture of SC with BO or BT treated with H_2_O, 10 µM (RXR)_4_XB-*acpP* targeting the *acpP* of SC or the respective scrambled control. OD600 data are shown as the mean of two independent experiments, and error bars indicate standard deviation. Illustrations created with BioRender. **c**,16S rRNA abundance relative to the H_2_O control of each species after treatment with 10 µM (RXR)_4_XB-scr or SC (RXR)_4_XB-*acpP*. Data from three independent starting cultures are shown; error bars represent standard deviation. The statistical significance of differences between treatments of each species was determined using two-way ANOVA with Bonferroni’s multiple comparisons test; ns, not significant; **** *P*⍰<⍰0.0001.

To monitor potential changes in the growth kinetics of *Bacteroides* upon *S. copri* depletion, we used a plate-based co-culture system that connects two individual wells with a semipermeable membrane, allowing diffusion of small molecules but not bacteria (Jo et al. 2023). Growth measurements of each bacterium demonstrate that ASO-mediated inhibition of *S. copri* growth does not affect the growth kinetics of *B. ovatus* or *B. thetaiotaomicron* individually over 24 hours (Fig. 5b). Next, to demonstrate that ASOs can selectively deplete a target species from an artificial community, we mixed the three species in equal parts and treated the resulting Bacteroidales community with *S. copri* (RXR)_4_XB-*acpP* as well as the respective scrambled ASO sequence control for 16 hours in order to allow all species to reach the stationary phase. The treatment with *S. copri* (RXR)_4_XB-*acpP* selectively depleted *S. copri* 100-fold and did not affect the abundance of *B. ovatus* or *B. thetaiotaomicron* (Fig. 5c). These results show that ASOs can be applied to selectively modulate a microbiota community without affecting members that are not targeted by the ASO.

## DISCUSSION

ASO-mediated translational inhibition of essential mRNAs has been established for multiple bacterial pathogens (El-Fateh et al. 2024), and recent studies suggest that ASOs may have applications beyond bacterial growth inhibition (Yokoi et al. 2022; Gerovac et al. 2025; Kawabuchi et al. 2025). Yet, to leverage their full potential as non-genetic tools for functional microbiota studies, it would be beneficial to extend ASO application to obligate anaerobic species and to investigate their efficacy in microbiota communities. Here, using multiple members of the *Prevotellaceae* family, we demonstrate efficient CPP-mediated ASO delivery, robust phenotypic readouts of translation inhibition in genetically intractable strains, and selective depletion of species from a community of anaerobic bacteria. To the best of our knowledge, this work is also the first to demonstrate true ASO multiplexing in bacteria, which will be important for dissecting relationships between functionally related genes in the absence of a good genetic system.

Due to their size and charge, ASOs do not passively cross the bacterial envelope (Xue et al. 2018; Ghosh et al. 2024), and thus need a carrier for efficient delivery to the cytoplasm. In line with previous reports showing that (RXR)_4_XB and (KFF)_3_K can efficiently deliver ASOs to the cytoplasm of Gram-negative bacteria (El-Fateh et al. 2024; Moreira et al. 2024; Pals et al. 2024), we found that these CPPs mediate ASO-dependent growth inhibition through translation inhibition of the essential gene *acpP* in *Prevotellaceae*. While (RXR)_4_XB-ASOs are thought to pass the bacterial envelope passively, (KFF)_3_K-ASOs rely on the inner membrane transporter SbmA (Yavari et al. 2021; Siekierska et al. 2024). Interestingly, *S. copri* lacks a SbmA homolog, suggesting either a transporter-independent uptake mechanism for (KFF)_3_K-ASOs or the presence of an alternative inner membrane transporter. When testing unconjugated CPPs for unspecific growth inhibition, we observed no effect of (RXR)_4_XB, but found that (KFF)_3_K showed growth delay at 20 µM and an MIC of 40 µM. This MIC value is consistent with other studies reporting an unconjugated (KFF)_3_K MIC of 32 µM for *E. coli* K12 and *L. monocytogenes*, as well as cytotoxic effects against *S. enterica* at 10 µM (Macyszyn et al. 2023; Ghosh et al. 2024). Therefore, out of eight CPPs tested, (RXR)_4_XB is the carrier of choice for *S. copri* based on efficient ASO delivery coupled with the low inherent cytotoxicity of the CPP itself.

ASO sensitivity varies between bacterial species due to differences in uptake efficiency and intrinsic resistance mechanisms (Pals et al. 2024; Moustafa et al. 2025). We found that *Prevotellaceae* species differed in MIC by more than 10-fold. The most sensitive species to ASO-mediated *acpP* inhibition was *P. corporis*, with an MIC of 2.5 µM, while the most resistant strain was *S. copri* HDC01, with an MIC of 40 µM. Interestingly, *S. copri* HDC01 was also the most resistant *Segatella* strain to genetic manipulation using a conjugation-based gene insertion system (Li et al. 2021). Generally, differences in MIC can be attributed to intrinsic properties of the ASO conjugates used or to strain diversification that affects membrane composition or biofilm formation (Xiao et al. 2024). The application of the same ASO to nine different strains from five different species yielded MICs ranging from 5 to 40 µM. The sensitivity was not associated with a specific *Segatella* species, as MIC values within *S. copri* strains ranged from 5 to 40 µM. These differences in sensitivity might also arise from some *Segatella* strains possibly possessing additional *acpP* genes, as observed with *S. hominis* HDD12. Indeed, out of the two strains displaying an MIC of 20 µM, *S. sinensis* HDE04 has a second predicted *acpP* gene that resembles *acpP*_2 of HDD12. Taken together, we observe that in *Prevotellaceae*, ASO efficiency depends on both the ASO-conjugate itself and the bacterial strain tested. Consequently, multiple ASO conjugates, ideally coupled with a CPP screen, should be evaluated when establishing ASOs for a given strain or species.

While growth inhibition is commonly used to assess ASO efficiency (Moustafa et al. 2025), phenotype-based readouts may be more specific, especially when unique morphological changes such as filamentation (*ftsZ*; Goh et al. 2009) or loss of mid-cell localization (*phuZ*; Gerovac et al. 2025) are induced. The targeting of *ftsZ* or *mreB* as used here should be easily transferable to other bacteria, since FtsZ and MreB are often conserved across species and protein knockdown has been linked to morphological changes in most bacteria studied (Battaje et al. 2023; Wang et al. 2025). However, these phenotypes might not be as pronounced in other species, and certain bacteria, such as *Chlamydia* spp. lack one of the two proteins (Gaballah et al. 2011). In addition, multiple gene copies, as we found in *S. copri* DSM 18205^T^, may obscure the expected phenotypes for single ASO treatments. Nonetheless, we expect *ftsZ-* and *mreB-*induced phenotypic changes to provide a robust read-out for targeted ASO inhibition in other microbiota species.

This study demonstrates a true multiplex application of ASOs, achieving simultaneous repression of up to three genes in a single cell. This should be of interest to microbiologists working with bacteria that lack robust forward genetics tools, where knocking out a single gene is time-consuming, let alone disrupting multiple genes. A multiplex tunable ASO application, as demonstrated for the essential genes *acpP* and *ftsZ*, facilitates the study of gene epistasis and synthetic lethality by enabling simultaneous translation inhibition of multiple mRNAs in a single cell (Klobucar and Brown 2018; González-González et al. 2024). Moreover, combinatorial ASO therapies could improve the treatment efficacy of antibacterial ASOs by targeting multiple genes to create synergy, as well as by simultaneously targeting mRNAs involved in resistance mechanisms and essential genes to inhibit growth more effectively. Thus, we anticipate that ASO multiplexing will be a powerful approach for advanced gene-function studies and antibacterial applications. In particular, we expect ASO multiplexing to be valuable for Bacteroidales species to dissect which polysaccharide utilization loci might cooperate to break down complex sugars, which is important for gut community studies and prebiotic development (Grondin et al. 2017; Qu et al. 2025).

Manipulation of microbiota communities is an emerging concept for targeted interventions and studying bacterial interactions (Cullen et al. 2020). As proof of concept that ASOs can be used to manipulate community composition, we showed selective inhibition of *S. copri* within a synthetic Bacteroidales community. This paves the way for further applications of ASO-based *in situ* manipulation of communities, enabling a more sophisticated dissection of interactions by repressing genes of interest in individual microbes and elucidating molecular mechanisms that affect community function. This could be especially interesting to study bacterial species that cannot be grown in monoculture and necessitate community settings for robust growth (D’Onofrio et al. 2010; Stewart 2012; Lloyd et al. 2018). Furthermore, community manipulation, such as the deletion or modification of detrimental species, could improve microbiota community properties for health preservation (Sheth et al. 2016). *S. copri* has been associated with numerous beneficial and detrimental impacts on human health, which are connected to strain-specific effects (Claus 2019; Amend et al. 2023; Nii et al. 2023). This points to the importance of targeted manipulation at the strain level as a potential therapeutic application. While studies on *S. copri* biology have been limited due to genetic inaccessibility, recent studies have identified molecular factors important for gut colonization, which could serve as ASO targets for programmable microbiota manipulation (El Mouali et al. 2024; Tawk et al. 2025).

Regarding the proof-of-concept community experiments conducted in this study, the observed selectivity for *Segatella* warrants further investigation. For example, we do not know the efficiency of CPP-mediated ASO uptake in *B. ovatus* and *B. thetaiotaomicron*, as it is difficult to predict whether *S. copri acpP* ASO was effectively delivered into the bacterial cytoplasm of the two *Bacteroides* species. In addition, gene essentiality might change upon co-culture conditions, especially for genes involved in metabolic pathways (Ibberson et al. 2017). So even though gene essentiality has been inferred from genetic screens using monoculture experiments, unexpected effects might be observed when ASOs are used in community settings. Nonetheless, since an ASO targeting *S. copri acpP* selectively depleted *S. copri* without affecting the other two *Bacteroides* species, ASOs have the potential for targeted manipulation of the microbiota.

Several factors need to be considered in extending this approach to other microbiota species. Many anaerobic bacteria are difficult to culture, and their slow growth might complicate the establishment of robust growth assays (Hitch et al. 2021). Thus, we established phenotypic readouts that allow for a clear observation of ASO-induced phenotypes, which should also be applicable to slow-growing bacteria. Furthermore, ASO applications might also be constrained to specific culture media that need to be adjusted to accommodate bacterial growth and efficient ASO delivery (Wornell et al. 2022; Kumar and Srivastava 2011; Choi et al. 2014). Here, we established ASOs targeting *Prevotellaceae* using YCFA medium, which enables rapid bacterial growth and high ASO efficiency. Importantly, YCFA allows robust growth of multiple anaerobic microbiota species, making it a good candidate for ASO screens in other bacterial species of the microbiota (Tidjani Alou et al. 2020).

In conclusion, our findings support ASOs as a strategy for selectively modulating anaerobic bacteria and for analyzing genetically intractable members of the gut microbiome. We envision ASOs as an easy-to-use, robust, and versatile technology for bacterial functional genomics and precision microbiome editing, poised to accelerate research into the human microbiota.

## Author Contributions

Conceptualization: V.C., P.K.D., L.P., T.S., and J.V. Methodology: V.C., P.K.D., Y.E.M., T.W., L.P., F.F., T.S., and J.V. Investigation: V.C., V.L., T.W., and V.G. Visualization: V.C. Writing the original draft: V.C., T.W., and J.V. Reviewing and editing the manuscript: V.C., T.W., L.P., F.F., T.S., and J.V.

## Data availability

All source data will be provided with this paper. The manuscript does not include data with mandatory deposition.

## Acknowledgements

We are grateful to Anke Sparmann for her exceptional help with editing the manuscript and the Vogel laboratory for great discussions. Additionally, we would like to thank Anna Zweyer and Luise Klos for their technical assistance. We thank the Vogel Stiftung Dr. Eckernkamp for supporting V.C. and V.L. with a Dr. Eckernkamp Fellowship. This work was supported by funds to J.V. from a DFG Gottfried Wilhelm Leibniz Award (DFG Vo875-18) and the Bavarian bayresq.net project Rbiotics (F.F., J.V.). Research was further funded by the BMBF in the framework of the Cluster4Future program (Cluster for Nucleic Acid Therapeutics Munich, CNATM; Project ID: 03ZU1201CA; L.P., J.V.), the German Research Foundation SFB 1583/1 (Project number: 492620490, Subproject A09; J.V.), the German Research Foundation SPP 2474 (Project number 564359880; F.F., J.V.), and the German Research Foundation - EXC 2155 (Project number 390874280, T.S.). F.F. and J.V. are members of the DFG Cluster of Excellence, Project-ID 533767322 – EXC 3113/1: Cluster for Nucleic Acid Sciences and Technologies (NUCLEATE).

## MATERIAL AND METHODS

### Bacterial strains and culture conditions

The following bacterial strains were used in this study: *Segatella copri* (formerly *Prevotella copri*) DSM 18205^T^, *Bacteroides thetaiotaomicron* DSM 2079^T^, *Bacteroides ovatus* DSM 1896^T^, *Prevotella corporis* DSM 18810^T^, and *Prevotella intermedia* DSM 20706^T^ acquired from the German Collection of Microorganisms and Cell Culture (DSMZ); *S. copri* DSM 108556 (HDC01), *S. copri* DSM 1085578 (HDD04), *S. copri* (RPC01), *S. copri* (RPA01), *S. brasiliensis* DSM 112105 (HDD05), *S*. brunsvicensis DSM 112023 (NI025), *S. hominis* DSM 111806 (HDD12), *S. sanihominis* DSM 113786 (HDD12), *S. sinensis* DSM 108151 (HDE04), *S. sinica* DSM 111807 (HDE06) were obtained from Till Strowig (Helmholtz Centre for Infection Research, Braunschweig, Germany). All strains were routinely grown in 80:10:10 (N_2_:H_2_:CO_2_) anaerobic conditions inside a Coy Vinyl anaerobic chamber (Coy Laboratory Products, Inc., USA) on supplemented BHI agar plates (brain–heart infusion (BHI), 2% agar, 1% (w:v) dried yeast extract, 1% (v:v) sterile filtered glucose solution, 5⍰µg⍰per ml of hemin, 10% (v:v) fetal bovine serum, 1 µg per ml Vitamin K_3_) for three days at 37°C from 20% glycerol stocks kept at -80°C. For liquid growth all strains were cultured in self-made degassed YCFA medium (modified from DMSZ 1611: Gibco™ Bacto™ Casitone (10 g/L), dried yeast extract (2.5 g/L), glucose (5.0 g/L), maltose (2.0 g/L), cellobiose (2.0 g/L), NaHCO_3_ (4 g/L), L-cysteine HCl·H_2_O (1 g/L), K_2_HPO_4_ (0.45 g/L), KH_2_PO_4_ (0.45 g/L), (NH_4_)_2_SO_4_ (0.9g/L), NaCl (0.9 g/L), MgSO_4_·7H_2_O (0.045 g/L), CaCl2·2H_2_O (0.045 g/L), resazurin (0.001 g/L), acetic acid (1.9 ml/L), propionic acid (0.6 ml/L), iso-butyric acid (0.1 ml/L), n-valeric acid (0.1 ml/L), iso-valeric acid (0.1 ml/L), hemin (0.01 g/L), biotin (0.002 g/L), Vitamin B_12_ (0.1 mg/L), folic acid (0.002 g/L), pyridoxine-HCl (0.01 g/L), p-aminobenzoic acid (0.005 g/L) and distilled H_2_O up to 1 L volume; pH was adjusted to 7.3) at 37°C inside the anaerobic chamber. Precultures were prepared 24 h before inoculating the working culture at a 1:50 dilution in YCFA until an OD600 of 0.4 (mid-exponential phase) was reached. Cultures were then diluted to the desired CFU/ml. All tubes, media, plates, and reagents were placed into the anaerobic chamber 24 h before use to ensure complete oxygen depletion.

### Cell-penetrating peptides (CPPs) and antisense oligomers (ASOs)

CPPs, ASOs, and CPP-ASO constructs were obtained from Peps4LS GmbH (Heidelberg), and all compounds were analyzed by mass spectrometry and HPLC to assess their quality and quantity (Table 1 and Supplementary Table 1). The ASO modality used in this study is peptide nucleic acid (PNA), a synthetic DNA analog with a peptide-like backbone. ASO sequences were designed with the help of MASON (Jung et al. 2023), and scrambled ASO and non-targeting control sequences were verified to have no off-targets in the translation initiation regions of the genomes of the targeted bacterial species by manual searches. CPP and ASO stocks were stored at -20°C, low-binding tips and low-binding tubes (Sarstedt) were used throughout for handling. To ensure solubility and correct stock preparation, all compounds were vortexed for three seconds, spun down, heated at 55°C for five minutes, and then again vortexed and centrifuged. Working stocks were prepared in ultra-pure water, and concentrations were determined using Nano-Drop spectrophotometer measurements at A_205nm_ for CPPs or A_260nm_ for CPP-ASOs, and adjusted if necessary.

### Minimum inhibitory concentration (MIC) determination and growth inhibition assays

To determine the MIC value or monitor growth kinetics, broth microdilutions were performed. A 24 h bacterial preculture was diluted in YCFA medium and grown to an OD600 of 0.4 (mid-exponential phase). The bacteria were then diluted 1:2500 to obtain 1 x 10^5^ CFU/ml. 95 µl of diluted bacterial cultures were pipetted into a transparent 96-well plate (Nunc™, Thermo Fischer Scientific) together with 5 µl of 20x CPP stocks, ASO stocks, or water as a control. Bacterial growth was monitored at 37°C by measuring OD600 every 20 minutes with constant shaking in a plate reader (Epoch 2, Biotek) positioned in the anaerobic chamber. The MIC was defined as the lowest concentration at which growth was visibly inhibited (OD600 <0.1).

### Determination of bactericidal effect using spotting

To investigate bactericidal effects of ASOs, a 24 h bacterial preculture grown in YCFA was diluted 1:50 in fresh YCFA and grown to an OD600 of 0.4. This culture was then diluted 1:2500 to 1 x 10^5^ CFU/ml in YCFA. 95 μl of the culture was dispensed into a transparent 96-well plate (Nunc™, Thermo Fisher Scientific) together with 5 μl of 20x ASO stock or water, and the plate was incubated at 37°C with shaking at 237 rpm every 20 minutes. At the sampled time points, 50 μl were removed from the well, a 1:10 serial dilution series was prepared with 450 μl anoxic 1x PBS, and 5 μl of each dilution were spotted onto anoxic, supplemented BHI plates for CFU determination. Plates were incubated for two to three days at 37°C in the anaerobic chamber before CFU counting and imaging.

### Microscopy

Bacterial phenotypes were investigated using confocal laser scanning microscopy (CLSM). Briefly, mid-exponential cultures were diluted to 10^5^ CFU/mL in YCFA and incubated with ASOs at 37°C and 230 rpm in the anaerobic chamber for 24 h. The samples were then removed from the chamber, centrifuged at 4°C for 10 minutes at 13,000 g to collect the pellet, fixed with 4% (w/v) PFA at 4°C for 10 minutes, washed once with 1x PBS, and resuspended in 1x PBS to an OD600 of approximately 1. Afterward, 1-2 μl of cell suspension was spotted on an 1.5% agar pad for CLSM imaging. Samples were imaged using ibidi 8-well chambers (ibidi) on a Leica Stellaris laser-scanning confocal microscope (Leica Microsystems) in bright-field mode. CLSM images were analyzed using ImageJ.

### Sequence identity determination and structure predictions

Amino acid identity of proteins was determined using Clustal Omega (1.2.4) multiple sequence alignment tool (Madeira et al. 2024). The structure of each protein was predicted using AlphaFold 3 (Abramson et al. 2024).

### Synthetic community treatment and quantitative PCR

For the synthetic community pre-cultures of *S. copri* DSM 18205^T^, *Bacteroides ovatus* DSM 1896^T^, and *Bacteroides thetaiotaomicron* DSM 2079^T^ were cultured in YCFA and incubated at 37°C for 24 h. The pre-cultures were then diluted 1:50 for *S. copri* or 1:500 for *B. ovatus* and BT and grown until an OD600 of 0.4. These cultures were diluted to 1 x 10^5^ CFU/ml and mixed in equal ratios for a total volume of 950 ml. Afterward, 50 µl of H2O or 20x ASO solution was added to the culture for a final concentration of 10 µM, and the community was incubated at 37°C with 230 rpm in the anaerobic chamber. After 16 h of treatment, the relative abundance of *S. copri, B. ovatus*, and *Bacteroides thetaiotaomicron* was monitored by quantitative PCR (qPCR) using strain-specific primers and a general bacterial 16S primer pair as a control. The sequences and sources of all primers used are listed in Supplementary Table 2. Genomic DNA was extracted from the bacterial community using chloroform extraction, and qPCR was performed on a CFX96 instrument (BioRad) with Takyon™ master mix (Eurogentec) and each primer at a final concentration of 100 nM. Cycling conditions were 95 °C for 10 minutes, followed by 50 cycles of 95 °C for 15 seconds and 60 °C for 60 seconds. To determine the efficiency of each primer pair, calibration curves were generated using a 10-fold serial dilution of a pool of all DNA extracts. Primer efficiencies were then used to calculate the abundance of 16S DNA for each strain relative to the total bacterial 16S rRNA content, using the Pfaffl method (Pfaffl 2001) (Supplementary Fig. 1).

### Plate-based co-culture

To investigate bacterial growth kinetics in co-culture, co-culture plates kindly provided by Prof. Dr. Rolf Kümmerli and Lukas Schwyter from the University of Zurich were used. Briefly, pre-cultures of *S. copri, B. ovatus*, and *B. thetaiotaomicron* were prepared in YCFA, diluted 1:50 for *S. copri* or 1:500 for *B. ovatus* and *B. thetaiotaomicron*, and grown until an OD600 of 0.4. The strains were then diluted to 1 x 10^5^ CFU/ml, and 95 µl of bacterial culture were pipetted into each corresponding well. The left well of each connected pair was filled with *S. copri*, and the right well was filled with either YCFA, *B. ovatus*, or *B. thetaiotaomicron*. To each well, either 5 µl of H2O, 20x stock of ASO targeting the *acpP* of *S. copri*, or the scrambled ASO control was added for a final concentration of 10 µM. Bacterial growth in co-culture at 37°C was monitored by measuring OD600 every 20 minutes with constant shaking in a plate reader (Epoch 2, Biotek) positioned in the anaerobic chamber.

### Statistical Analysis

All statistical tests were done in GraphPad Prism (10.5.0, 2025) using two-way ANOVA with Bonferroni’s multiple comparisons test. A P value equal to or less than 0.05 was considered statistically significant.

## Extended Data

**Extended Data Fig. 1:**
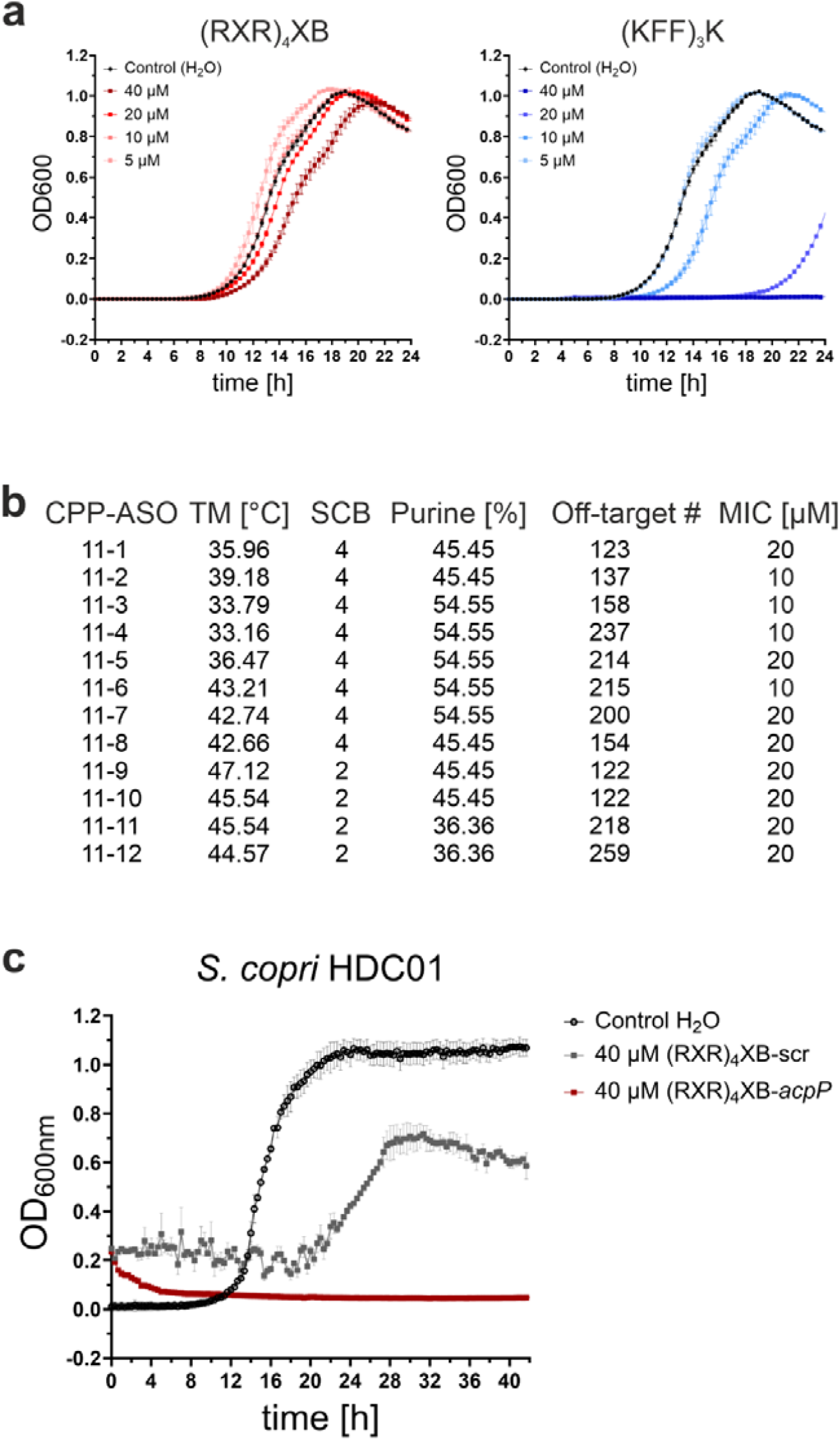
Characterization of ASO application within S. copri. **a**, Growth kinetics of *S. copri* DSM 18205^T^ treated with (RXR)_4_XB or (KFF)_3_K CPPs. Data are shown as the mean of two independent experiments; error bars indicate standard deviation; MIC indicated in bold. **b**, Melting temperature (TM), maximum stretch of self-complementarity bases (SCB), percentage of purine content, and off-target number of tiling CPP-ASOs according to MASON. Respective MIC value shown on the right. **c**, Growth kinetics of *S. copri* HDC01 treated with H_2_O, 40 µM of (RXR)_4_XB-*acpP* or the respective (RXR)_4_XB-scr control. OD600 data are shown as the mean from two independent cultures shown. Error bars indicate standard deviation.

**Extended Data Fig. 2:**
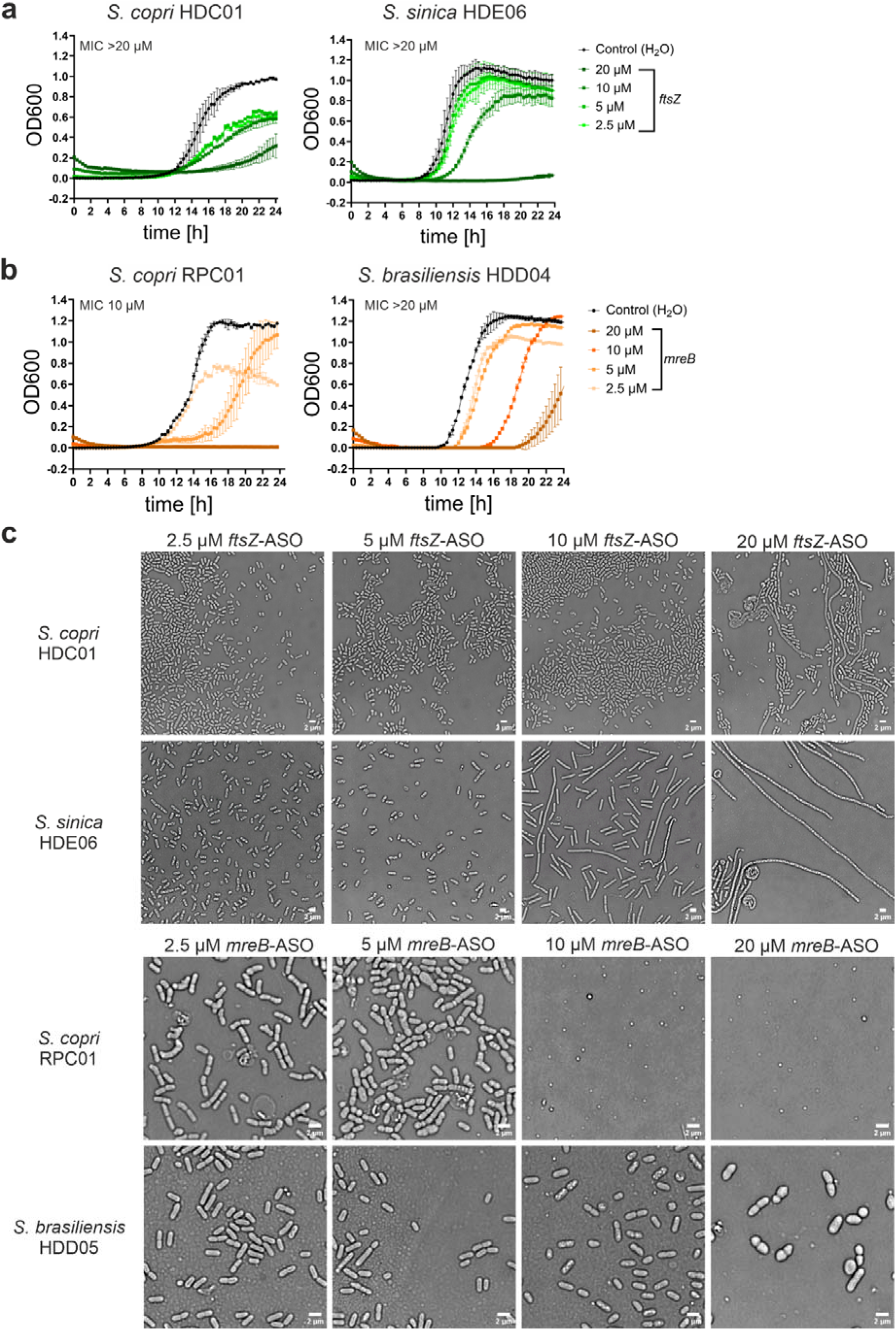
MIC determination of ftsZ- and mreB-targeting ASOs together with phenotype microscopy images across four genetically intractable strains of the S. copri complex. **a**, Growth kinetics of *S. copri* HDC01 and *S. sinica* HDE06 treated with H_2_O or titration series of (RXR)_4_XB-*ftsZ*. OD600 data are shown as the mean of two independent cultures. Error bars indicate standard deviation. MIC is indicated. **b**, Growth kinetics of *S. copri* RPC01 and *S. brasiliensis* HDD05 treated with H_2_O or titration series of (RXR)_4_XB-*mreB*. OD600 data are shown as the mean of two independent cultures. Error bars indicate standard deviation. MIC is indicated. **c**, Representative bright field images from two independent experiments of *S. copri* HDC01 and *S. sinica* HDE06 treated with a titration series of (RXR)_4_XB-ASOs targeting *ftsZ* (top) or *S. copri* RPC01 and *S. brasiliensis* HDD05 treated with a titration series of (RXR)_4_XB-ASOs targeting *mreB* (bottom). Scale bar, 2 µm.

**Extended Data Fig. 3:**
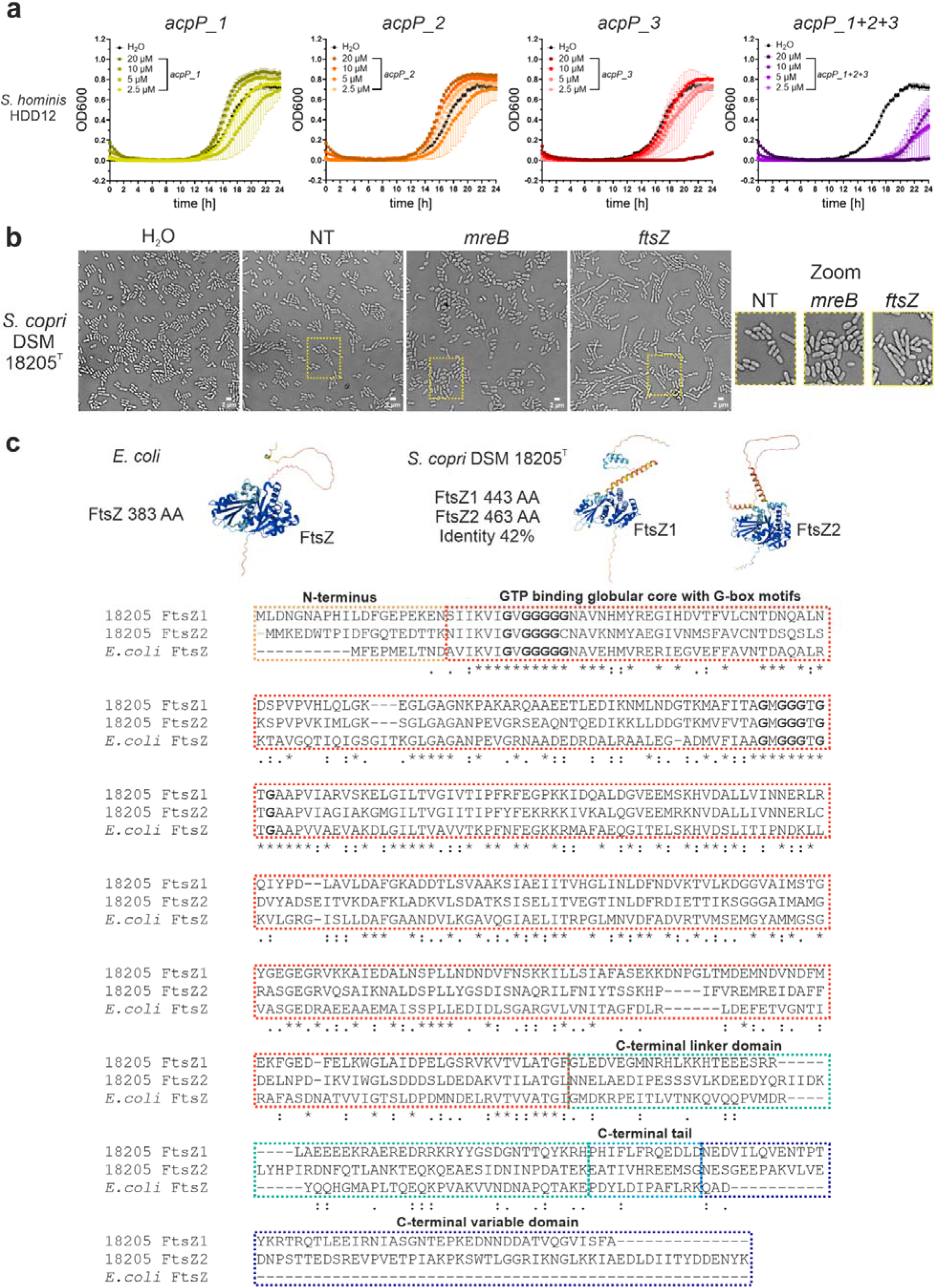
Multiplex ASO targeting enables the investigation of essential gene homologs within the S. copri complex. **a**, Growth curves of *S. hominis* HDD12 upon treatment with the indicated ASOs. OD600 data are shown as the mean of three independent starting cultures with standard deviation. **b**, Representative bright field images from two independent experiments of *S. copri* 18205^T^ treated with H_2_O, 10 µM of non-targeting ASO control, 10 µM (RXR)4XB-*mreB*, or 10 µM (RXR)_4_XB-*ftsZ*. Scale bar, 2 µm. **c**, (Top) FtsZ structures modeled by AlphaFold 3 from *E. coli* as well as *S. copri* 18205^T^. (Bottom) Protein sequence alignment performed by Clustal Omega for *E. coli* FtsZ, *S. copri* 18205^T^ FtsZ1, and *S. copri* 18205^T^ FtsZ2. Protein domains are indicated. Glycine residues from G-box motifs are highlighted in bold.

**Extended Data Fig. 4:**
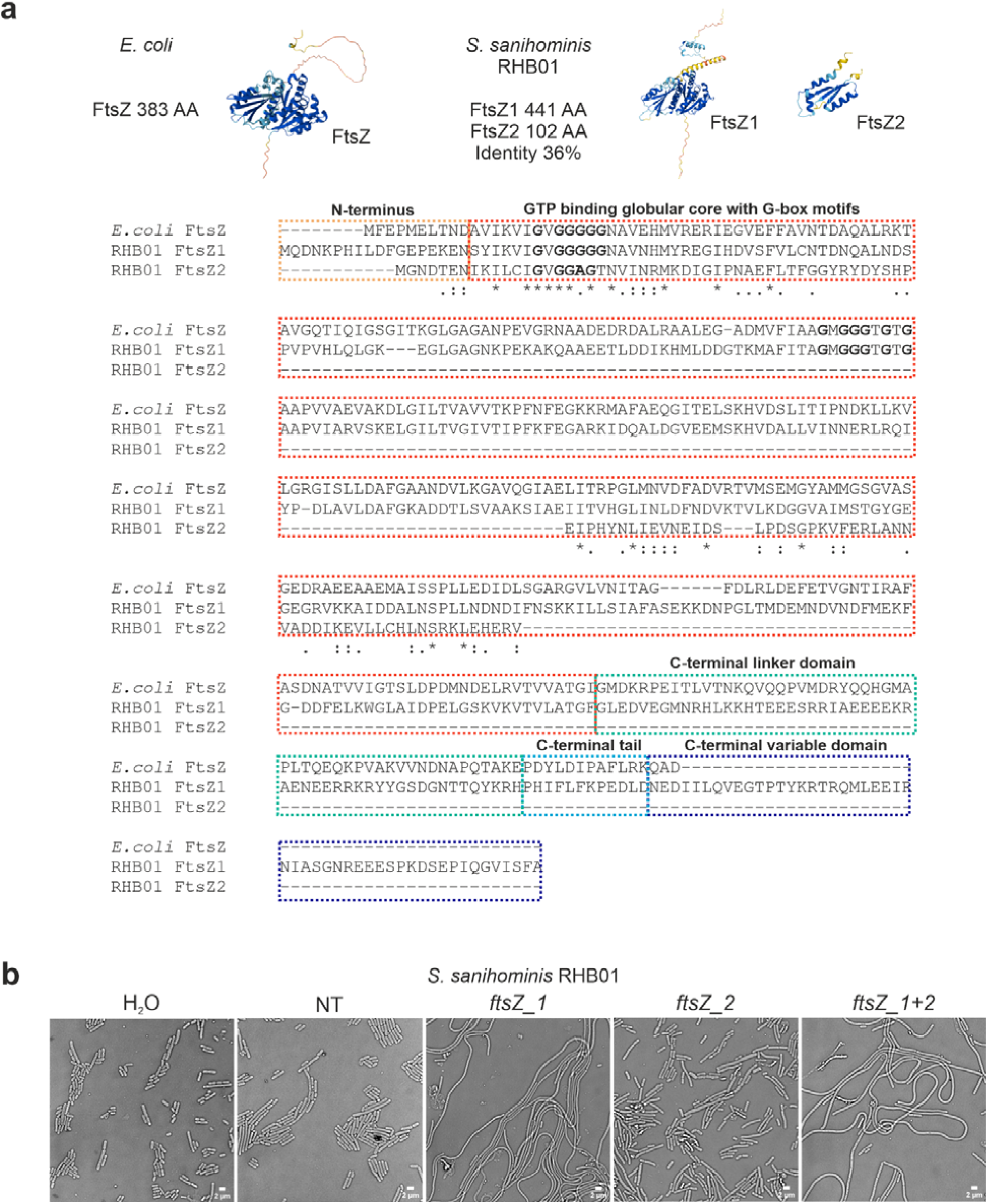
Multiplex ASO targeting reveals diversification of FtsZ homolog function in S. sanihominis RHB01. **a**, (Top) FtsZ structures modeled by AlphaFold 3 from *E. coli* as well as *S. sanihominis* RHB01. (Bottom) Protein sequence alignment performed by Clustal Omega for *E. coli* FtsZ, *S. sanihominis* RHB01 FtsZ1, and *S. sanihominis* RHB01 FtsZ2. Protein domains are indicated. Glycine residues from G-box motifs are highlighted in bold. **b**, Representative bright field images of *S. sanihominis* RHB01 from two independent experiments, treated with H_2_O, 5 µM non-targeting control (NT), or *ftsZ* (RXR)_4_XB-ASOs. Scale bar, 2 µm.

