## Supplementary Information for "ASO-mediated mRNA silencing enables functional analysis and selective depletion of the human microbiota *Prevotellaceae*"

### Supplementary Figure and Tables

Valentina Cosi[1], Vincent Lau[1,2], Petia Kovatcheva-Datchary[2], Youssef El Mouali[3], Toby Wilkinson[2], Victoria Gebler[2], Linda Popella[2,4], Franziska Faber[1,5,6], Till Strowig[3,7,8], Jörg Vogel[1,2,4,6]*

[1] Helmholtz Institute for RNA-based Infection Research (HIRI), Helmholtz Centre for Infection Research (HZI), 97080, Würzburg, Germany

[2] University of Würzburg, Medical Faculty, Institute of Molecular Infection Biology (IMIB), RNA Biology Group, 97080, Würzburg, Germany

[3] Helmholtz Center for Infection Research (HZI), 38142, Braunschweig, Germany

[4] Cluster for Nucleic Acid Therapeutics Munich (CNATM), Munich, Germany

[5] University of Würzburg, Medical Faculty, Institute for Hygiene and Microbiology, 97080, Würzburg, Germany

[6] Cluster for Nucleic Acid Sciences and Technologies – NUCLEATE

[7] Cluster of Excellence RESIST (EXC 2155), Hannover Medical School, 30625 Hannover, Germany

[8] Center for Individualized Infection Medicine (CiiM), a joint venture of Hannover Medical School and Helmholtz Centre for Infection Research, Hannover, Germany

Keywords: *Prevotellaceae*, cell-penetrating peptide, antisense oligomer, non-genetic tool, microbiota modulation

**This file includes:** Supplementary Figure 1 and Supplementary Tables.

Supplementary Figure 1.

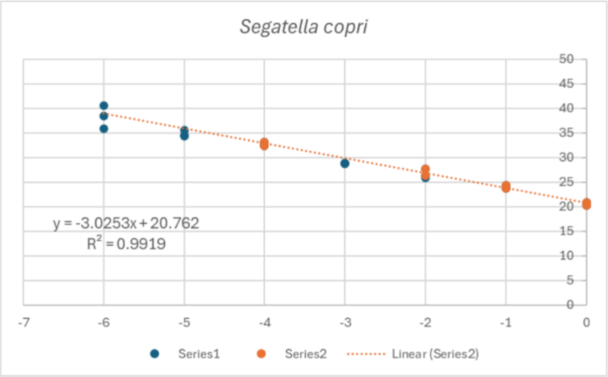


| | |
|---|---|
| Slope | -3.03 |
| Intercept | 20.76 |
| E | 2.14 |
| Efficiency | 1.14 |

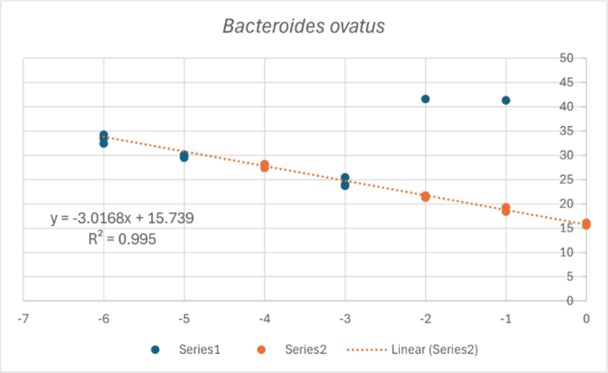


| | |
|---|---|
| Slope | -3.01 |
| Intercept | 15.74 |
| E | 2.15 |
| Efficiency | 1.15 |

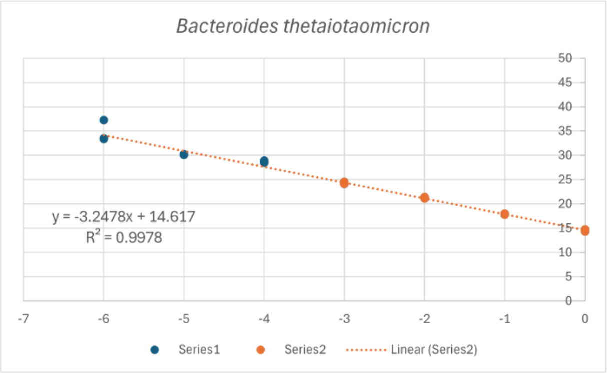


| | |
|---|---|
| Slope | -3.24 |
| Intercept | 14.62 |
| E | 2.03 |
| Efficiency | 1.03 |

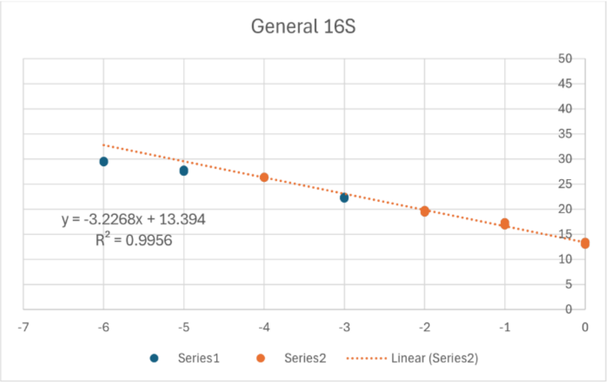


| | |
|---|---|
| Slope | -3.26 |
| Intercept | 13.39 |
| E | 2.04 |
| Efficiency | 1.04 |

***Supplementary Fig. 1: Calibration curves of qPCR primers from S. copri, B. ovatus, B. thetaiotaomicron, and general bacteria 16S.***

*Curves were created using a 10-fold serial dilution of a pool of all DNA extracts. Series 1 represents all observed data points. Series 2 represents the data points chosen from the linear range for efficiency calculations according to the MIQE 2.0 guidelines. All standard curves comprise four data points within the serial dilution series. Primer efficiencies (E – 1, E = 10-1/slope) used for sample calculations according to the Pfaffl method are depicted on the right of each calibration curve.*

Supplementary Table 1: Antisense oligomer conjugates

| | Internal # | Name | Target | Peptide sequence* | ASO sequence[#] | Length | Source |
|---|---|---|---|---|---|---|---|
| 1 | 78 | $(KFF)_3K$-*acpP* | *acpP* | KFFKFFKFFK | GACATAATTGT | 11mer | Peps4LS |
| 2 | 152 | $(KFF)_3K$-*acpP*-scr | - | KFFKFFKFFK | TAGACTATATG | 11mer | Peps4LS |
| 3 | 79 | $(RXR)_4XB$-*acpP (06)* | *acpP* | RXRRXRRXRRXRXB | GACATAATTGT | 11mer | Peps4LS |
| 4 | 153 | $(RXR)_4XB$-*acpP*-scr | - | RXRRXRRXRRXRXB | TAGACTATATG | 11mer | Peps4LS |
| 5 | 1981 | K8-*acpP* | *acpP* | KKKKKKKK | GACATAATTGT | 11mer | Peps4LS |
| 6 | 1984 | K8-*acpP*-scr | - | KKKKKKKK | TAGACTATATG | 11mer | Peps4LS |
| 7 | 2115 | R8-*acpP* | *acpP* | RRRRRRRR | GACATAATTGT | 11mer | Peps4LS |
| 8 | 2118 | R8-*acpP*-scr | - | RRRRRRRR | TAGACTATATG | 11mer | Peps4LS |
| 9 | 2116 | Seq35-*acpP* | *acpP* | RKKRRQRR | GACATAATTGT | 11mer | Peps4LS |
| 10 | 2119 | Seq35-*acpP*-scr | - | RKKRRQRR | TAGACTATATG | 11mer | Peps4LS |
| 11 | 2117 | Seq2483-*acpP* | *acpP* | HKDRKRWWLRGKKKKRKKKQK | GACATAATTGT | 11mer | Peps4LS |
| 12 | 2120 | Seq2483-*acpP*-scr | - | HKDRKRWWLRGKKKKRKKKQK | TAGACTATATG | 11mer | Peps4LS |
| 13 | 1982 | Tat-*acpP* | *acpP* | GRKKRRQRRRYK | GACATAATTGT | 11mer | Peps4LS |
| 14 | 1985 | Tat-*acpP*-scr | - | GRKKRRQRRRYK | TAGACTATATG | 11mer | Peps4LS |
| 15 | 1980 | $(RFR)_4XB$-*acpP* | *acpP* | RFRRFRRFRRFRXB | GACATAATTGT | 11mer | Peps4LS |
| 16 | 1983 | $(RFR)_4XB$ -*acpP*-scr | - | RFRRFRRFRRFRXB | TAGACTATATG | 11mer | Peps4LS |
| 17 | 2410 | $(RXR)_4XB$-*acpP_01* | *acpP* | RXRRXRRXRRXRXB | AATTGTAACTT | 11mer | Peps4LS |
| 18 | 2411 | $(RXR)_4XB$-*acpP_02* | *acpP* | RXRRXRRXRRXRXB | TAATTGTAACT | 11mer | Peps4LS |
| 19 | 2412 | $(RXR)_4XB$-*acpP_03* | *acpP* | RXRRXRRXRRXRXB | ATAATTGTAAC | 11mer | Peps4LS |
| 20 | 2413 | $(RXR)_4XB$-*acpP_04* | *acpP* | RXRRXRRXRRXRXB | CATAATTGTAA | 11mer | Peps4LS |
| 21 | 2414 | $(RXR)_4XB$-*acpP_05* | *acpP* | RXRRXRRXRRXRXB | ACATAATTGTA | 11mer | Peps4LS |
| 22 | 2415 | $(RXR)_4XB$-*acpP_07* | *acpP* | RXRRXRRXRRXRXB | TGACATAATTG | 11mer | Peps4LS |
| 23 | 2416 | $(RXR)_4XB$-*acpP_08* | *acpP* | RXRRXRRXRRXRXB | CTGACATAATT | 11mer | Peps4LS |
| 24 | 2417 | $(RXR)_4XB$-*acpP_09* | *acpP* | RXRRXRRXRRXRXB | TCTGACATAAT | 11mer | Peps4LS |
| 25 | 2418 | $(RXR)_4XB$-*acpP_10* | *acpP* | RXRRXRRXRRXRXB | TTCTGACATAA | 11mer | Peps4LS |
| 26 | 2419 | $(RXR)_4XB$-*acpP_11* | *acpP* | RXRRXRRXRRXRXB | TTTCTGACATA | 11mer | Peps4LS |
| 27 | 2420 | $(RXR)_4XB$-*acpP_12* | *acpP* | RXRRXRRXRRXRXB | ATTTCTGACAT | 11mer | Peps4LS |
| 28 | 2240 | $(RXR)_4XB$-*ftsZ*_S. copri | *ftsZ* | RXRRXRRXRRXRXB | TAGCATAGTCG | 11mer | Peps4LS |
| 29 | 2428 | $(RXR)_4XB$-*ftsZ*_HDE06 | *ftsZ* | RXRRXRRXRRXRXB | TAGCATAATAG | 11mer | Peps4LS |

| | | | | | | | |
|---|---|---|---|---|---|---|---|
| 30 | 2156 | $(RXR)_4$XB-*mreB_S. copri* | *mreB* | RXRRXRRXRRXRXB | CATTTCGTTTC | 11mer | Peps4LS |
| 31 | 2426 | $(RXR)_4$XB-*acpP1*_HDD12 | *acpP1* | RXRRXRRXRRXRXB | TCCATGTGATA | 11mer | Peps4LS |
| 32 | 2754 | $(RXR)_4$XB-*acpP2*_HDD12 | *acpP2* | RXRRXRRXRRXRXB | TCCATACGATT | 11mer | Peps4LS |
| 33 | 2756 | $(RXR)_4$XB-*acpP3*_HDD12 | *acpP3* | RXRRXRRXRRXRXB | GACATAATTTT | 11mer | Peps4LS |
| 34 | 2160 | $(RXR)_4$XB-*ftsZ2_S. copri* | *ftsZ2* | RXRRXRRXRRXRXB | TCATCATTTCT | 11mer | Peps4LS |
| 35 | 2509 | $(RXR)_4$XB-*mreB*_HDD05 | *mreB* | RXRRXRRXRRXRXB | CATTTGTTGTA | 11mer | Peps4LS |
| 36 | 2837 | $(RXR)_4$XB-*acpP_P. corporis* | *acpP* | RXRRXRRXRRXRXB | GACATAATTTC | 11mer | Peps4LS |
| 37 | 2840 | $(RXR)_4$XB-*acpP_P. intermedia* | *acpP* | RXRRXRRXRRXRXB | TCTGACATAAT | 11mer | Peps4LS |

*Peptide sequences are shown from N to C terminus. #PNA sequences are shown from N to C terminus

### Supplementary Table 2: qPCR primers

| | Internal | Oligonucleotide sequence | Description | Source | Publication |
|---|---|---|---|---|---|
| 1 | 19563 | CGAAAGCTTGCTTTTGATGG | *S. copri* 16S forward primer | Sigma | Verbrugghe et al. 2021 |
| 2 | 19564 | CGCAAGGTTATCCCCAAGT | *S. copri* 16S reverse primer | Sigma | Verbrugghe et al. 2021 |
| 3 | 19801 | TGCAAACTRAAGATGGC* | *B. ovatus* 16S forward primer | Sigma | Tong et al. 2011 |
| 4 | 19802 | CAAACTAATGGAACGCATC | *B. ovatus* 16S reverse primer | Sigma | Tong et al. 2011 |
| 5 | 19797 | GCAAACTGGAGATGGCGA | *B. thetaiotaomicron* 16S forward primer | Sigma | Tong et al. 2011 |
| 6 | 19798 | AAGGTTTGGTGAGCCGTTA | *B. thetaiotaomicron* 16S reverse primer | Sigma | Tong et al. 2011 |
| 7 | 19567 | GTGSTGCAYGGYYGTCGTCA* | General bacteria 16S forward primer | Sigma | Rinttilä et al. 2004 |
| 8 | 19568 | ACGTCRTCCMCNCCTTCCTC* | General bacteria 16S reverse primer | Sigma | Rinttilä et al. 2004 |

*R = purine, Y = pyrimidine, S = guanine/cytosine, M = adenine/cytosine, N = adenine/guanine/cytosine/thymine

Reference list primers:

Rinttilä, T., A. Kassinen, E. Malinen, L. Krogius, and A. Palva. 2004. "Development of an Extensive Set of 16S rDNA - targeted Primers for Quantification of Pathogenic and Indigenous Bacteria in Faecal Samples by Real - time PCR." Journal of Applied Microbiology 97 (6): 1166–77. https://doi.org/10.1111/j.1365-2672.2004.02409.x.

Tong, Jia, Chengxu Liu, Paula Summanen, Huaxi Xu, and Sydney M. Finegold. 2011. "Application of Quantitative Real-Time PCR for Rapid Identification of Bacteroides Fragilis Group and Related Organisms in Human Wound Samples." Anaerobe 17 (2): 64–68. https://doi.org/10.1016/j.anaerobe.2011.03.004.

Verbrugghe, Phebe, Olivier Van Aken, Frida Hållenius, and Anne Nilsson. 2021. "Development of a Real-Time Quantitative PCR Method for Detection and Quantification of Prevotella Copri." BMC Microbiology 21 (1): 23. https://doi.org/10.1186/s1
